# Neural correlates of extinction in a rat model of appetitive Pavlovian conditioning

**DOI:** 10.1101/2022.09.28.509892

**Authors:** Alexa Brown, Franz R. Villaruel, Nadia Chaudhri

**Author notes:** Corresponding Author: Alexa Brown, B.A., Center for Studies in Behavioural Neurobiology, Department of Psychology, Concordia University, 7141 Sherbrooke Street West, Montreal, QC, H4B-1R6, Canada.

## Abstract

Extinction is a fundamental form of inhibitory learning that is important for adapting to changing environmental contingencies. While numerous studies have investigated the neural correlates of extinction using Pavlovian fear conditioning and appetitive operant reward-seeking procedures, less is known about the neural circuitry mediating the extinction of appetitive Pavlovian conditioned responding. Here, we aimed to generate an extensive brain activation map of extinction learning in a rat model of appetitive Pavlovian conditioning. Male Long-Evans rats were trained to associate a conditioned stimulus (CS; 20 s white noise) with the delivery of a 10% sucrose unconditioned stimulus (US; 0.3 ml/CS) to a fluid port. Control groups also received CS presentations, but sucrose was delivered either during the inter-trial interval or in the home-cage. After conditioning, 1 or 6 extinction sessions were conducted in which the CS was presented but sucrose was withheld. We performed Fos immunohistochemistry and network connectivity analyses on a set of cortical, striatal, thalamic, and amygdalar brain regions. Neural activity in the prelimbic cortex, ventral orbitofrontal cortex, nucleus accumbens core, and paraventricular nucleus of the thalamus was greater during recall relative to extinction. Conversely, prolonged extinction following 6 sessions induced increased neural activity in the infralimbic cortex, medial orbitofrontal cortex, and nucleus accumbens shell compared to home-cage controls. All these structures were similarly recruited during recall on the first extinction session. These findings provide novel evidence for the contribution of brain areas and neural networks that are differentially involved in the recall versus extinction of appetitive Pavlovian conditioned responding.

## Introduction

Through associative learning, animals and humans can adapt their behaviour in response to changes in their environment in order to survive. In Pavlovian conditioning, organisms learn to associate a conditioned stimulus (CS; e.g., white noise) with an unconditioned stimulus (US; e.g., sucrose). As a result, the CS comes to elicit conditioned responding. In contrast, in operant conditioning, organisms learn to perform an operant response (e.g., lever press) to receive an outcome (e.g., sucrose). During extinction, responding can be reduced by repeated exposure to the CS alone and withholding the outcome. However, even after extensive extinction training, the memory formed during conditioning is retained and can provide a powerful basis for relapse (Bouton, 2002). Instead of erasing the original learning, extinction is thought to involve new brain plasticity that encodes a “CS-no US” or “response-no outcome” association (Bouton et al., 2006). Conditioning- and extinction-related memories are thought to be largely separate from each other (Bouton & Swartzentruber, 1991; Bouton, 2002; Lacagnina et al., 2019). The neurobiology mediating these distinct learning and memory processes has been investigated using Pavlovian fear conditioning and appetitive operant conditioning procedures (Maren, 2001; Milad & Quirk, 2002; Corcoran & Maren, 2004; Peters et al., 2008; Marchant et al., 2010; LaLumiere et al., 2012; Moorman et al., 2015; Warren et al., 2016; 2019). However, appetitive Pavlovian procedures have primarily been used to investigate the neural mechanisms of associative learning (Saddoris et al., 2009; Keefer & Petrovich, 2017), and considerably less is known about the brain areas and networks that mediate the extinction of appetitive Pavlovian conditioned responding.

The infralimbic cortex (IL) is thought to be a critical brain area mediating the extinction of Pavlovian fear conditioning and operant reward-seeking (Quirk et al., 2000; Milad & Quirk, 2002; Peters et al., 2009). In support of this hypothesis, IL neurons are active during fear extinction retrieval, and during appetitive operant extinction learning and expression (Knapska & Maren, 2009; Marchant et al., 2010; Orsini et al., 2011, 2013; Warren et al., 2016). Moreover, activation of the IL with the glutamate receptor agonist AMPA, or with the AMPA potentiator PEPA, suppresses the reinstatement of operant reward-seeking, suggesting that enhanced excitatory synaptic activity in the IL inhibits operant reward-seeking (Peters et al., 2008; LaLumiere et al., 2012). Similarly, stimulation of the IL enhances fear extinction learning and retrieval (Milad et al., 2004; Vidal-Gonzalaz et al., 2006; Adhikari et al., 2015; Do-Monte et al., 2015a). Consistent with a role of the IL in inhibiting responding, inactivation of the IL disinhibits operant reward-seeking and promotes reinstatement (Peters et al., 2008; Gutman et al., 2017). Consistently, inhibition of the IL impairs fear extinction memory consolidation and potentiates freezing (Burgos-Robles et al., 2007; Laurent and Westbrook, 2008; Sotres-Bayon et al., 2009; Sierra-Mercado et al., 2011; Do-Monte et al., 2015a; Lebrón et al., 2004; Farrell et al., 2010). Together, these results support a role of the IL in mediating the extinction of operant reward-seeking and Pavlovian fear conditioning.

A functional dichotomy in the medial prefrontal cortex (mPFC) has been suggested in which the IL mediates the extinction of conditioned responding, while the prelimbic cortex (PL) drives the expression of responding (Peters et al., 2009). The PL is thought to drive the expression of conditioned fear and operant reward-seeking through downstream projections to the basolateral amygdala (BLA) and nucleus accumbens core (NAcC), respectively (Peters et al., 2009). While the neural pathways from the IL to the BLA and nucleus accumbens shell (NAcSh) are thought to mediate the extinction of Pavlovian fear conditioning and operant reward-seeking, respectively (Peters et al., 2009). Support for this hypothesis comes from the findings that PL inactivation disrupts the expression of conditioned fear but does not affect extinction learning or retrieval (Laurent & Westbrook, 2009; Choi et al., 2010; Sierra-Mercado et al., 2011). Moreover, Fos expression is increased in PL inputs to the BLA during the retrieval of fear conditioned memories, while inhibition of the PL-to-BLA pathway impairs the retrieval of fear conditioned memories (Do-Monte et al., 2015b; Quiñones-Laracuente et al., 2021). Similarly, inhibition of the PL-to-NAcC pathway attenuates the reinstatement of operant cocaine-seeking (Stefanik et al., 2013, 2016). Conversely, chemogenetic activation of the IL-to-NAcSh pathway suppresses the reinstatement of operant reward-seeking, while simultaneous inhibition of the IL and NAcSh drives reinstatement (Peters et al., 2008; Augur et al., 2016). Likewise, activation of IL inputs to the amygdala enhances fear extinction learning and retrieval, while inhibition of the IL-to-BLA pathway impairs fear extinction learning and retrieval (Adhikari et al., 2015; Bukalo et al., 2015; Bloodgood et al., 2018). Together, these results support a functional distinction within the mPFC such that the IL and PL mediate the extinction and expression of conditioning responding, respectively, and these functions are maintained through distinct projections to the BLA and NAc based on the valence of the reinforcer.

There is also evidence that the BLA, orbitofrontal cortex (OFC), and paraventricular nucleus of the thalamus (PVT) are implicated in motivated behaviours and may be involved in both the expression and extinction of conditioned responding. Inactivation of the BLA impairs the extinction of operant food-seeking (McLaughlin & Floresco, 2007). Similarly, inactivation of the OFC disrupts extinction learning in both rats and non-human primates (Butter et al., 1963; Panayi & Killcross, 2014). Moreover, single-unit recordings have shown that the BLA represents learning in over-expectation, and this learning appears to be supported via inputs from the OFC (Lucantonio et al., 2015). In over-expectation, a decline in Pavlovian conditioned responding, like that observed in extinction, is produced by the over-expectation, rather than the omission of the outcome. Therefore, both the BLA and OFC appear to be implicated in behavioural inhibition produced by both extinction and over-expectation. However, inactivation of the BLA, or the OFC-to-BLA pathway, also disrupts the renewal of extinguished drug-seeking, suggesting that the OFC and BLA may also be implicated in driving the expression of conditioned responding for rewarding outcomes (Fuchs et al., 2005, 2007; Marinelli et al., 2010; Lasseter et al., 2011). More recent evidence has also implicated the PVT in the expression and extinction of conditioned responding via distinct projections from the PL and IL, respectively. Inhibition of the PL-to-PVT pathway attenuates the reinstatement of extinguished cocaine-seeking (Giannotti et al., 2018), while inhibition of the IL-to-PVT pathway impairs fear extinction retrieval (Tao et al., 2021), suggesting that the IL/PL functional dichotomy is maintained through projections to the PVT.

Although few studies have assessed whether the neural mechanisms of extinction in operant reward-seeking studies extend to appetitive Pavlovian learning models, the IL also appears to mediate the extinction of appetitive Pavlovian conditioned responding. Optogenetic stimulation of the IL suppresses context-induced renewal of appetitive Pavlovian conditioned responding after extinction (Villaruel et al., 2018). In addition, lesions of the IL disrupt extinction retrieval, and promote the reinstatement and renewal of appetitive Pavlovian conditioned responding after extinction (Rhodes & Killcross, 2004; 2007). These results parallel findings from appetitive operant conditioning studies and support a role of the IL in mediating appetitive Pavlovian extinction. In contrast, however, inactivation of the IL *facilitates* the extinction of appetitive Pavlovian conditioned responding (Mendoza et al., 2015). Moreover, recent evidence from our laboratory shows that optogenetic stimulation of IL inputs to the NAcSh attenuates renewal of appetitive Pavlovian conditioned responding, although *not* through an extinction mechanism (Villaruel et al., 2022). Together, these results suggest that the IL may mediate appetitive Pavlovian extinction, but the role of the NAcSh is unclear because of inconsistencies in the available evidence.

The goal of the current study was to examine the neural basis of appetitive Pavlovian extinction by examining the extent of activation in multiple brain regions during recall and following extinction. We examined a set of cortical, striatal, thalamic, and amygdalar brain regions, and assessed brain activation by measuring differences in Fos density, an immediate early gene widely used for brain activity mapping (Franceschini et al., 2020). Based on these data, we computed inter-regional correlations of Fos density to produce functional network activation graphs (Silva et al., 2019) for the recall and extinction of appetitive Pavlovian conditioned responding.

## Methods

### Animals

Subjects were 38 experimentally naïve, male Long-Evans rats (Charles River, St. Constant, Quebec, Canada; 220-240 g upon arrival). Rats were maintained in a humidity (40-45%) and climate controlled (21 °C) room on a 12-h light/dark cycle with lights turned on at 7:00 h. All procedures occurred during the light phase. Rats were individually housed in standard cages containing beta chip bedding (Aspen Sani chips; Envigo, Indianapolis IN) with unrestricted access to water and food (Agribands, Charles River). Each cage contained a nylabone toy (Nylabones; Bio-Serv, Flemington, NJ), a polycarbonate tunnel (Rat Retreats, Bio-Serv) and shredded paper. During a seven-day acclimation period to the colony room, rats were handled, and body weight was recorded daily. All procedures followed the guidelines of the Canadian Council for Animal Care and were approved by the Concordia University Animal Research Ethics Committee.

### Apparatus

Experiments were conducted in sound-attenuating melamine cubicles that each contained a conditioning chamber (Med Associates, ENV-009A, St-Albans, VT, USA). Each conditioning chamber consisted of stainless-steel bar floors, and a house light (75 W, 100 mA, ENV-215M) was in the center of the left wall. A white noise generator and speaker (ENV-225SM) in the top left corner of the left wall produced the white noise conditioned stimulus (CS) 5 dB above the background noise of an exhaust fan mounted inside the cubicle. A 10% sucrose solution was delivered to a fluid port (ENV-200R3AM) located 2 cm above the floor in the center of the right wall. Infrared sensors (ENV-254CB) lined both sides of the port opening to detect port entries. Solutions were delivered into the fluid port via a polyethylene tube (Fisher Scientific, 141 691 A) connected to a 20 ml syringe in a pump (Med Associates, PHM-100, 3.33 rpm) located outside the cubicle. All events were controlled and recorded by Med Associates software (Med-PC IV) on a computer in the experimental room.

### Behavioural procedures

Rats were acclimated to the taste of 10% sucrose in the home-cage for 48 h, and subsequently assigned into one of three experimental conditions: paired (n=14), unpaired (n=12) and home-cage (n=12). Groups were matched according to body weight, sucrose preference, and sucrose consumption. All rats were then habituated to the experimental chambers on one day (20 min session) during which the house light turned on after a 1 min delay and total port entries were recorded.

Rats were then trained in a Pavlovian conditioning task for 8 days (57 min sessions). During Pavlovian conditioning, the house-light turned on after a 2 min delay to signal the start of each session, and shut off to signal the end of the session. For the paired condition, Pavlovian conditioning sessions consisted of 10 pairings of an auditory CS (20 s white noise) that co-terminated with the delivery of 0.3 ml of 10% sucrose solution into a fluid port (10 s; 3 ml per session). The variable inter-trial interval (ITI) averaged 240 s (120, 240, or 360 s). Ports were checked after each session to ensure consumption.

Rats in the unpaired and home-cage conditions received identical sessions except that the sucrose was not delivered contingent to the CS. Rather, rats in the unpaired condition were presented with the same number of white noise presentations and sucrose deliveries as the paired condition, but sucrose was delivered to the fluid port mid-way during the ITI. Rats in the home-cage condition received the same number of white noise presentations as the paired and unpaired conditions and following the same ITI schedule, however, sucrose was delivered in the home-cage at a random time 1-4 h after the session. These control groups helped to determine whether Fos expression was specific to Pavlovian conditioned responding. In contrast to the home-cage control condition, where only the white noise stimulus was delivered, the CS and US were presented at intervals that prevented the establishment of an excitatory CS-US association for the unpaired control condition.

Following Pavlovian conditioning, rats were assigned to one of two experimental groups: recall (n=19; n=7 paired, n=6 unpaired, n=6 home-cage) and extinction (n=19; n=7 paired, n=6 unpaired, n=6 home-cage). Groups were matched according to ΔCS port entries (CS minus pre-CS port entries) during the training phase. Rats in the recall group received 1 extinction session (57 min session) prior to collection of tissue for Fos immunohistochemistry, while rats in the extinction group received 6 days of extinction training (57 min sessions). Extinction sessions were identical to Pavlovian conditioning sessions except that sucrose delivery was withheld.

### Fos immunohistochemistry and quantification

To quantify Fos expression induced by recall or extinction, rats were sacrificed 90 min after the start of either the first or sixth extinction session (Warren et al., 2016). Rats were anesthetized with sodium pentobarbital (100 mg/kg, intraperitoneal) and transcardially perfused with 0.1 M phosphate buffered saline (PBS), followed by 4% paraformaldehyde (PFA) in 0.1 M PBS. Brains were extracted and post fixed for 24 h in 4% PFA, before transfer to a 30% sucrose solution in water for 48 h. Coronal sections (40 μm) were obtained with a cryostat and collected in 0.1 M phosphate buffer (PB). Sections were blocked for 1 h in 0.3% PBS-Triton-X-100 with 6% normal goat serum (NGS; Vector Labs, S-1000), followed by incubation for 72 h at 4 ºC with anti-cFos rabbit primary antibody (1:2000; Cell Signalling, 2240). Sections were washed 3 × 10 min with PBS, and then incubated in a biotinylated goat anti-rabbit secondary antibody (1:250; Vector Labs, BA-1000) for 1 h in 0.3% PBS-Triton-X-100 and 3% NGS. Sections were washed 3 × 10 min with PBS and incubated in a tertiary of avidin and biotinylated horseradish peroxidase (1:1000; ABC kit, Vector Labs, PK-6100) and stained with a 3, 3’-diaminobenzidine (DAB) solution. Sections were washed in PB, mounted on slides and cover slipped.

Selected brain regions of interest (Table 1) were chosen a priori for statistical comparisons. A bright field microscope (Nikon Eclipse TiE) captured images at 20 × magnification. Images were imported into Fiji (ImageJ) software, and the number of Fos-positive neurons in each region of interest was quantified using a custom-made cell-counting Fiji macro, which quantified Fos-positive cells based on contrast with background, size, and circularity. A rat brain atlas (Paxinos and Watson, 2007) was used to approximate the location of sections compared to bregma. We quantified the number of Fos-positive neurons at the following bregma coordinates: 3.24, 3.00, 2.76 for the IL; 4.68, 4.20, 3.24 for the PL; 4.68, 4.20 for the mOFC; 4.20, 3.24 for the lOFC and vOFC; 2.28, 2.04, 1.20 for the NAcC and mNAcSh; 2.04, 1.20 for the lNAcSh; - 1.56, -1.80, -2.76, -3.24 for the BLA; -1.80, -2.04 for the aPVT; -2.64, -2.76 for the mPVT, and -3.00, -3.24 for the pPVT. The area used for quantification in all brain regions was selected manually and was consistent across all rats. All cells within that area were quantified for each section. Cell counts were then divided by the area selected in Fiji to calculate density. The density of Fos-positive neurons was averaged across sections for each rat. Density counts for the paired and unpaired conditions were normalized to the average density in the home-cage condition, which was used as a baseline (Barbosa et al., 2013; Lopez et al., 2018).

**Table 1.**
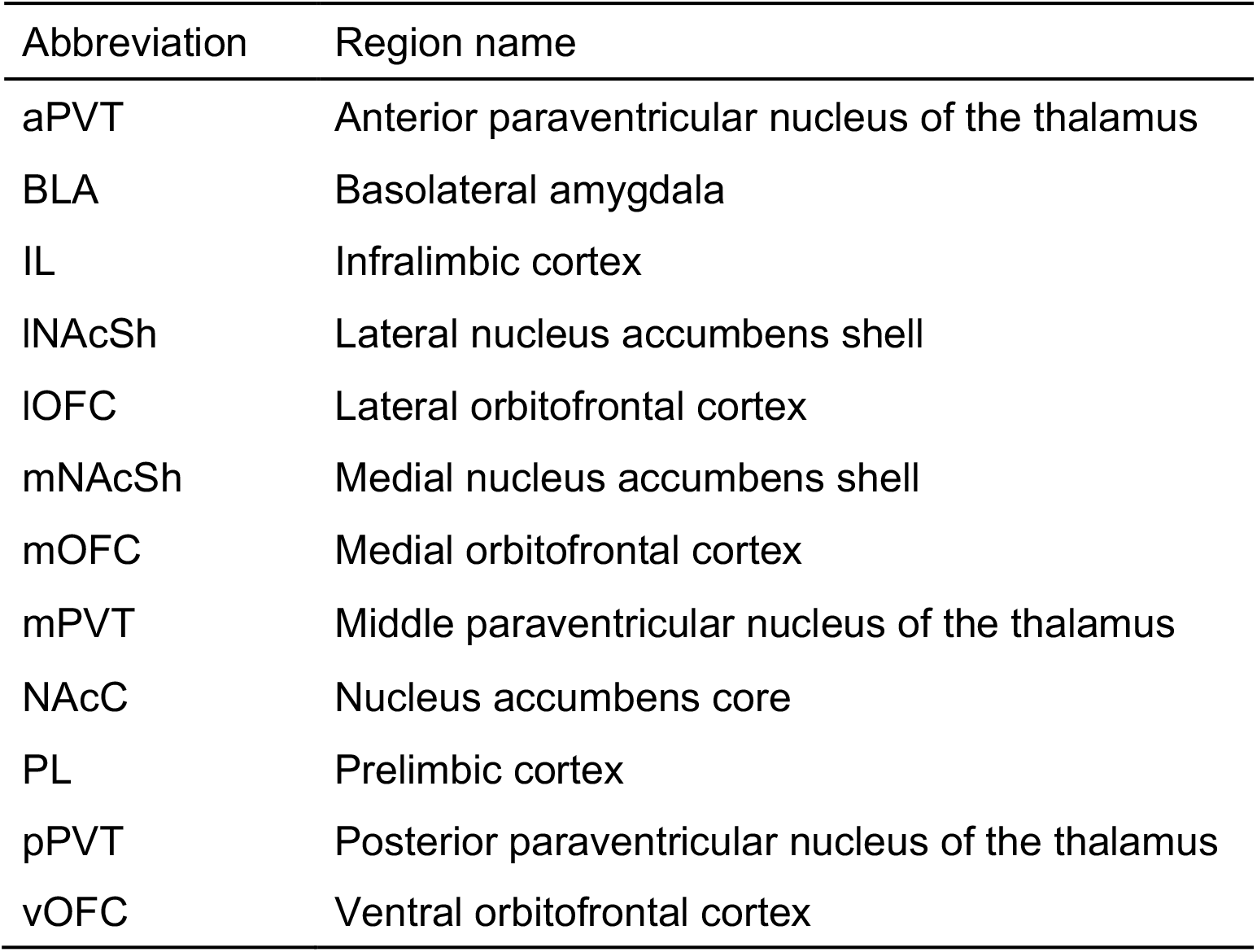
Abbreviations of brain areas

### Data analysis

The acquisition of ΔCS port entries (CS minus pre-CS port entries) was assessed using mixed analyses of variance (ANOVA) with group (recall or extinction), condition (paired, unpaired or home-cage), and session as factors. ΔCS port entries on the first extinction session were assessed using an ANOVA with group (recall or extinction) and condition (paired, unpaired or home-cage) as factors. Only in the extinction groups, extinction of ΔCS port entries was assessed using mixed ANOVA with condition (paired, unpaired or home-cage) and session as factors. ΔCS port entries during the Fos induction session (extinction session 1 or 6) were assessed using an ANOVA with group (recall or extinction) and condition (paired, unpaired or home-cage) as factors.

Fos density in each region of interest was analyzed separately using ANOVAs with group (recall or extinction) and condition (paired, unpaired or home-cage) as factors.

Greenhouse-Geisser corrections are reported following violations of Mauchly’s test of sphericity. Post-hoc analyses were corrected for multiple comparisons using the Bonferroni adjustment. All data analyses were conducted using IBM SPSS v21.0 (IBM Corp., Armonk, NY). Results were considered statistically significant at p<.05.

Network activation graphs were constructed using Fos density and Pearson correlation coefficients (Silva et al., 2019). Within each experimental group and condition (e.g., home-cage recall), Fos densities were averaged across bregma coordinates for each rat, thus each rat was an n=1 for each brain region. For all 12 brain regions analyzed, Pearson correlation coefficients were then calculated for Fos density between all pairwise combinations (Silva et al., 2019). Each node represents one of the 12 brain regions examined in this study. Node sizes in the paired and unpaired conditions are proportional to the Fos density increase for each brain region compared to the Fos densities in the corresponding home-cage control condition (Silva et al., 2019). Each edge connecting two nodes represents a Pearson correlation between brain regions that had a p<.05 and an r ≥.05 (adapted from Silva et al., 2019). Edge thickness reflects the r value of the correlation between the two brain regions, with thicker edges indicating a greater correlation in Fos densities in the two regions. Negative correlations are not represented in the network activation graphs. Correlations were displayed as a colour-coded correlation matrix using GraphPad Prism (Version 7; La Jolla, CA). The NetworkX (v2.7.1) package (Hagberg et al., 2008) in Python was used to visualize network activation graphs.

## Results

### Behavioural paradigm to assess Fos expression during recall and extinction

To assess brain activation patterns during recall and extinction of appetitive Pavlovian conditioned responding, we established a behavioural paradigm that could efficiently condition and extinguish responding to a Pavlovian CS (Fig. 1A). Rats in the paired condition received presentations of a 20 s white-noise CS that was paired with a 10% sucrose US, while rats in the unpaired and home-cage conditions received sucrose either during the ITI or in the home-cage after the session. Following training, rats in the recall groups received 1 extinction session, while rats in the extinction groups received 6 extinction sessions (Fig. 1B). Brains were extracted 90 minutes following the start of either the first or sixth extinction session and analyzed for Fos expression.

**Fig. 1.**
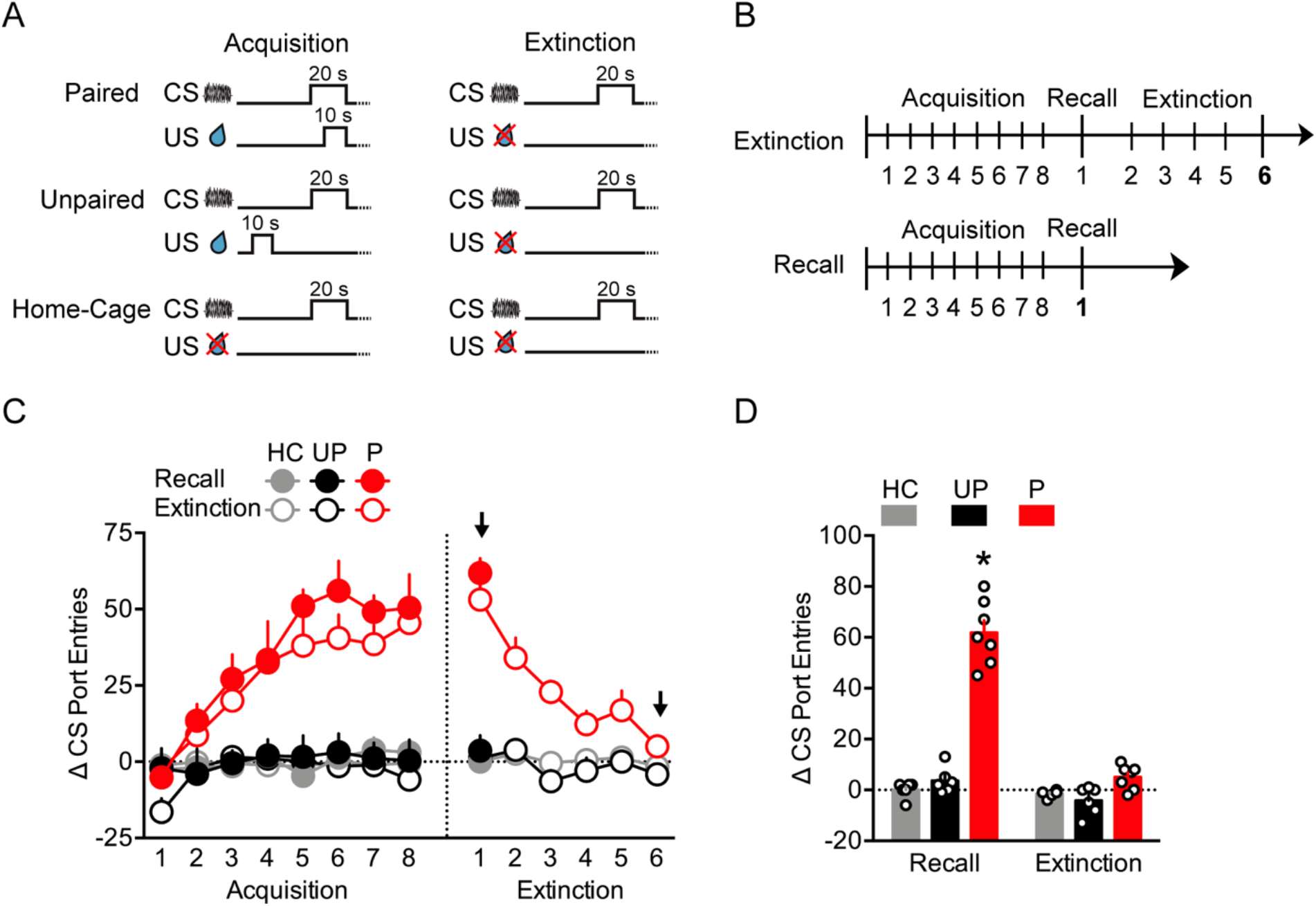
Acquisition and extinction of conditioned responding to a discrete sucrose CS. **A** Schematic representation of the experimental paradigm. During acquisition, the paired condition received pairings of a 20 s white noise CS that co-terminated with delivery of a sucrose US. The unpaired and home-cage conditions received white noise presentations that were not contingent with sucrose delivery and received sucrose either during the ITI or in the home-cage. During extinction, the white noise was presented as in acquisition, but sucrose was withheld. **B** Schematic diagram indicating the timing of acquisition and extinction sessions. Tissue was collected for Fos analysis 90 min after the first extinction session in the recall groups, and after the sixth extinction session in the extinction groups. **C** Changes in ΔCS port entries during acquisition show the development of Pavlovian conditioned responding only in the paired groups. Responding was maintained in both paired groups after one extinction session (recall) and responding was extinguished over the 6 extinction sessions in the paired extinction group. Arrows indicate sessions after which tissue was collected for Fos analysis. **D** ΔCS port entries collapsed across the two Fos induction sessions were increased for the paired recall group compared to the paired extinction, unpaired recall and home-cage recall groups. *p<.05, recall vs extinction in the paired condition. Here and in subsequent figures, data are depicted as mean ± SEM, and individual data points are overlaid on the bar graphs.

During conditioning, ΔCS port entries increased across sessions in the paired groups and remained low for the unpaired and home-cage groups (Fig. 1C; Condition, F_1,32_=60.77, p<.001; Session, F_3.5,112.2_=14.80, p<.001; Condition x Session, F_3.5,112.2_=9.38, p<.001). There was no significant difference in ΔCS port entries during conditioning between the recall and extinction groups that were formed by matching based on ΔCS port entries during conditioning (Group, F_1,32_=1.63, p=.211; Group x Condition x Session, F_3.5,112.2_=.61, p=.748).

During the first extinction session, ΔCS port entries were greater for the paired recall and paired extinction groups, compared to the the unpaired and home-cage conditions (Fig. 1C; Condition, F_2,32_=68.67, p<.001; Group, F_1,32_=.63, p=.434; Condition x Group, F_2,32_=.34, p=.718). Responding was significantly increased in the paired groups compared to the unpaired (p<.001) and home-cage (p<.001) groups with no difference between the unpaired and home-cage groups (p=1.000). Across the 6 extinction sessions, ΔCS port entries decreased for the paired extinction group, while the unpaired and home-cage extinction groups showed stable, low levels of responding (Fig. 1C; Condition, F_2,16_=30.39, p<.001; Session, F_2.7,44.0_=8.68, p<.001; Condition x Session, F_2.7,44.0_=5.73, p<.001).

Collapsed across both Fos induction sessions, ΔCS port entries were significantly greater for the paired recall group compared to all other groups (Fig. 1D; Condition, F_2,32_=113.86, p<.001; Group, F_1,32_=102.91, p<.001; Group x Condition, F_2,32_=68.02, p<.001). There were no significant differences between the paired extinction, unpaired extinction and home-cage extinction groups (all p>.05). However, responding was significantly increased in the paired recall group compared to the paired extinction group (p<.001). Therefore, only rats in the paired conditions learned to respond by entering the fluid port during the CS during conditioning, and only rats in the paired extinction group extinguished ΔCS port entries across extinction sessions.

### Fos immunoreactivity

#### Prelimbic and Infralimbic cortex

In the prelimbic cortex (PL), there was a main effect of condition, in which both the paired and unpaired groups showed markedly greater Fos density relative to the home-cage condition (Fig. 2A, B, C; Condition, F_2,32_=7.96, p=.002). Fos density did not differ significantly between the paired and unpaired conditions (p=1.000) but Fos density was greater in both the paired (p=.002) and unpaired (p=0.018) conditions relative to the home-cage control. There was also a significant main effect of recall vs extinction, in which Fos density was greater during recall as compared to extinction (Group, F_1,32_=5.33, p=.028), but there was no significant Group x Condition interaction (F_2,32_=1.30, p=.286), indicating that the lower Fos density in the extinction groups did not depend upon history of conditioning.

**Fig. 2.**
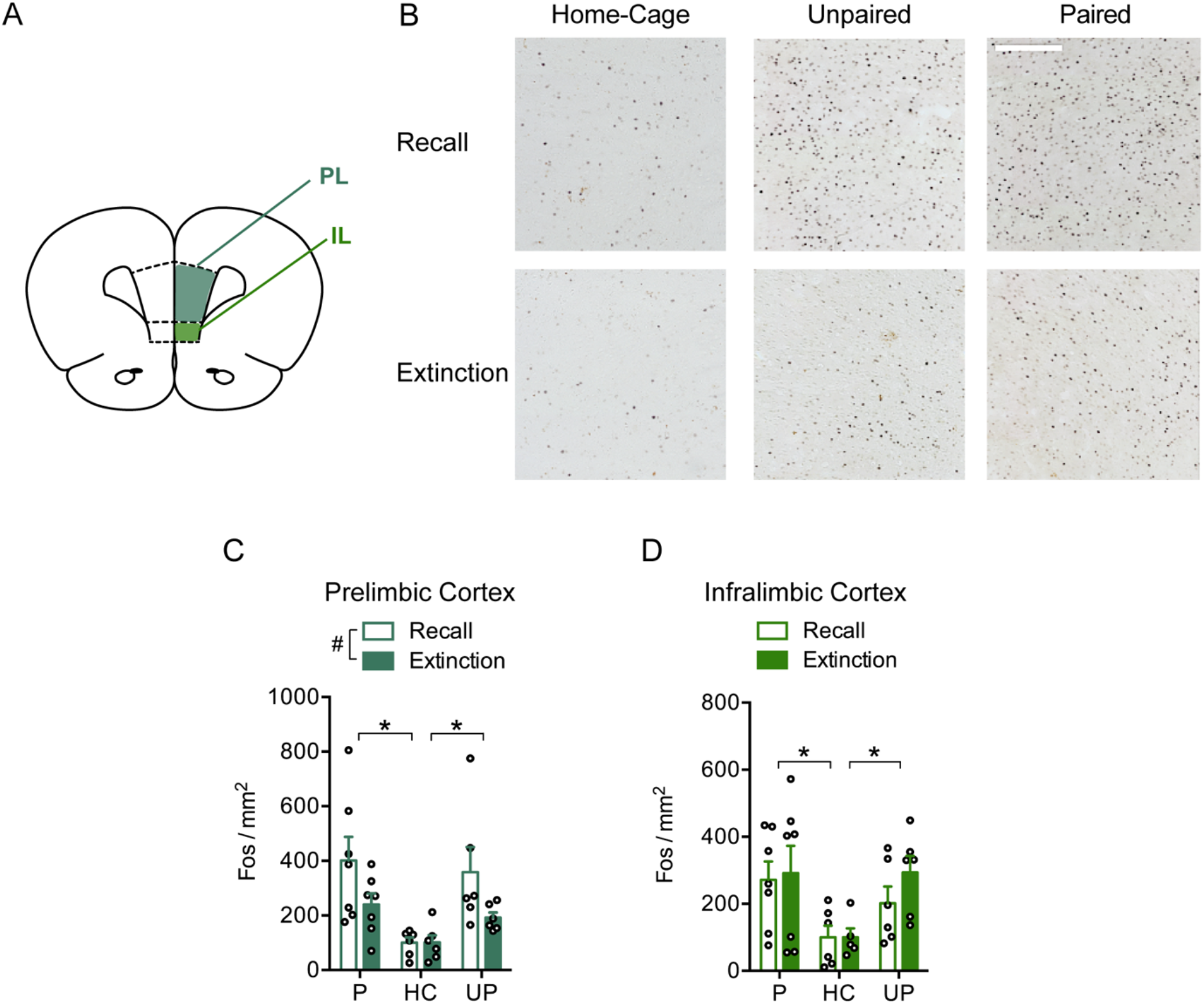
Fos density in the prelimbic (PL) and infralimbic (IL) cortices. **A** A schematic diagram indicates regions where Fos was quantified in the PL and IL. **B** Representative Fos photomicrographs in the PL. Scale bar is 250 μm. **C** Fos density in the PL was greater following recall relative to extinction, and for the paired and unpaired groups relative to home-cage groups. **D** Fos density in the IL was not different between recall and extinction and was greater for the paired and unpaired groups relative to home-cage groups. #p<.05, recall vs extinction main effect. *p<.05, post hoc comparisons following main effect of training condition.

In the infralimbic cortex (IL), Fos density was similar during recall and extinction (Fig. 2A, D; Group, F_1,31_=.66, p=.423). Similar to results for the PL, Fos density in the IL was markedly greater in the paired and unpaired groups relative to the home-cage groups (Condition, F_2,31_=5.68, p=.008). Fos density was not significantly different between the paired and unpaired conditions (p=1.000) but was greater in the paired (p=0.009) and unpaired (p=.049) conditions relative to the home-cage condition. There was no Group x Condition interaction (F_2,31_=.36, p=.702).

#### Orbitofrontal cortex

In the medial orbitofrontal cortex (mOFC), Fos density was greater in the paired and unpaired groups relative to the home-cage groups (Fig. 3A, B, C; Condition, F_2,31_=10.39, p<.001). Fos density was not significantly different between the paired and unpaired conditions (p=1.000) but was greater in the paired (p<.001) and unpaired (p=.005) conditions relative to the home-cage condition. Like results in the IL, Fos density was similar during recall and extinction (Group, F_1,31_=.02, p=.894), and there was no Group x Conditioned interaction (Group x Condition, F_2,31_=.50, p=.610).

**Fig. 3.**
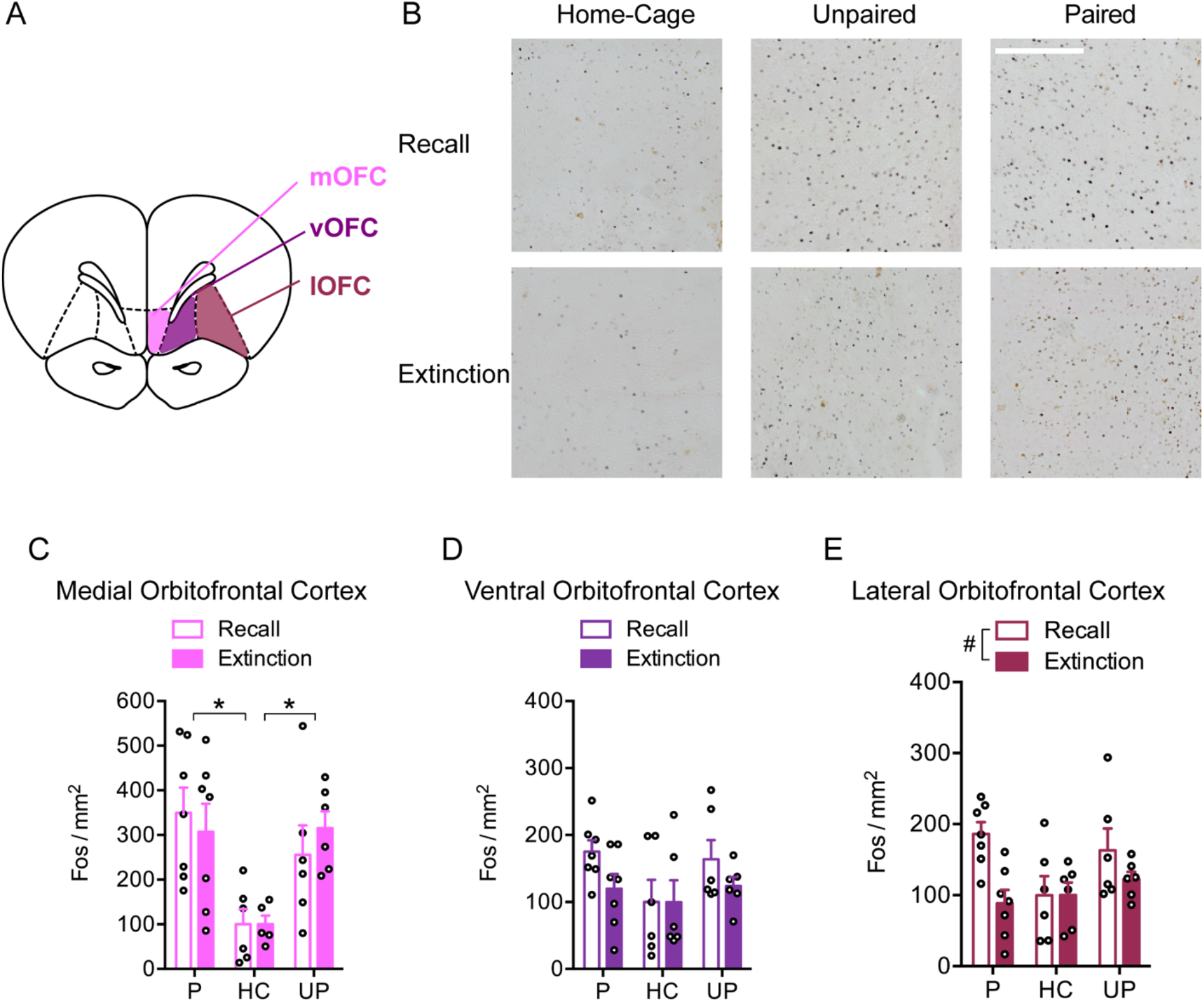
Fos density in the medial orbitofrontal cortex (mOFC), ventral orbitofrontal cortex (vOFC), and lateral orbitofrontal cortex (lOFC). **A** A schematic diagram indicates regions where Fos was quantified in the mOFC, vOFC, and lOFC. **B** Representative Fos photomicrographs in the mOFC. Scale bar is 250 μm. **C** Fos density in the mOFC was not different between recall and extinction, and was greater for the paired and unpaired conditions relative to the home-cage condition. **D** Fos density in the vOFC was not different between recall and extinction and was similar between the paired, unpaired, and home-cage conditions. **E** Fos density in the lOFC was greater following recall relative to extinction, and was similar between the paired, unpaired, and home-cage conditions. #p<.05, recall vs extinction main effect. *p<.05, post hoc comparisons following main effect of training condition.

In contrast to other cortical regions, in the ventral orbitofrontal cortex (vOFC), there was no significant difference in Fos density between the recall and extinction groups, and there were no significant differences between the paired, unpaired, and home-cage conditions (Fig. 3A, D; Group, F_1,32_=2.42, p=.130; Condition, F_2,32_=2.18, p=.129; Group x Condition, F_2,32_=.65, p=.527).

In the lateral orbitofrontal cortex (lOFC), there was no significant differences between paired, unpaired and home-cage conditions (Fig. 3A, E; Condition, F_2,32_=2.38, p=.109). There was a significant main effect of recall vs extinction, in which Fos density was greater during recall as compared to extinction (Group, F_1,32_=7.23, p=.011), but there was no significant Group x Condition interaction (F_2,32_=2.81, p=.075), indicating that the lower Fos density in the extinction groups did not depend upon history of conditioning.

#### Nucleus accumbens

In the nucleus accumbens core (NAcC), there was a main effect of condition in which the paired and unpaired groups showed greater Fos density compared to the home-cage controls (Fig. 4A, B, C; Condition, F_2,31_=10.86, p<.001). There was no significant difference between the paired and unpaired conditions (p=1.000), but Fos density was markedly greater in the paired (p=0.002) and unpaired (p=0.001) conditions relative to the home-cage condition. Like results in the PL, there was a main effect of recall vs extinction, in which Fos density was greater during recall compared to extinction (Group, F_1,31_=6.49, p=.016), but there was no Group x Condition interaction (F_2,32_=1.70, p=.199).

**Fig. 4.**
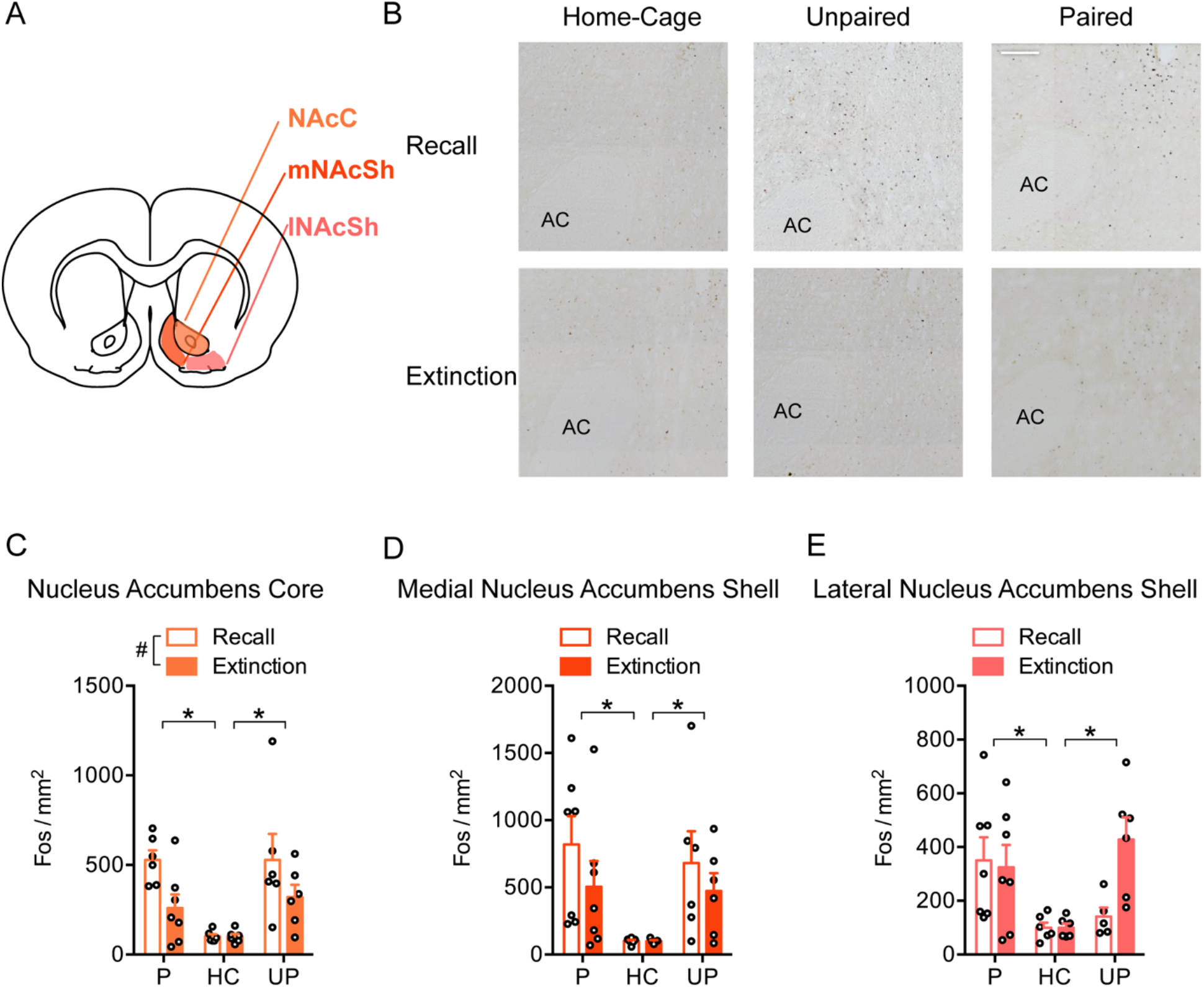
Fos density in the nucleus accumbens core (NAcC), medial nucleus accumbens shell (mNAcSh), and lateral nucleus accumbens shell (lNAcSh). **A** A schematic diagram indicates regions where Fos was quantified in the NAcC, mNAcSh, and lNAcSh. **B** Representative Fos photomicrographs in the NAcC. Scale bar is 200 μm. Anterior commissure (AC). **C** Fos density in the NAcC was greater following recall relative to extinction, and for the paired and unpaired conditions relative to the home-cage condition. **D** Fos density in the mNAcSh, and **E** lNAcSh, was not different between recall and extinction, and was greater for the paired and unpaired conditions relative to the home-cage condition. #p<.05, recall vs extinction main effect. *p<.05, post hoc comparisons following main effect of training condition.

Fos density in the medial and lateral nucleus accumbens shell (mNAcSh and lNAcSh), was found to be greater in the paired and unpaired conditions relative to the home-cage condition, but there was no difference between recall and extinction groups. In the mNAcSh, there was a main effect of condition (Fig. 4A, D; Condition, F_2,31_=5.88, p=.007) indicating that Fos density was no different between the paired and unpaired conditions (p=1.000) but was markedly greater in the paired (p=.008) and unpaired (p=.036) conditions relative to the home-cage controls. There was no significant difference between recall and extinction groups (Group, F_1,31_=1.53, p=.225) and no significant Group x Condition interaction (F_2,31_=.43, p=.658). Similarly, in the lNAcSh, Fos density was similar between the paired and unpaired conditions (p=1.000) but was greater in the paired (p=.003) and unpaired (p=.037) conditions relative to the home-cage condition (Fig. 4A, E; Condition, F_2,31_=7.10, p=.003). There was no significant main effect of recall vs extinction (Group, F_1,31_=2.52, p=.123), and no Group x Condition interaction (F_2,31_=3.19, p=.055).

#### Basolateral amygdala

In contrast to other brain regions, the basolateral amygdala (BLA) showed no significant differences in Fos density between the recall and extinction groups, as well as no differences between the paired, unpaired, and home-cage conditions (Fig. 5A, B, C; Group, F_1,32_=.17, p=.687; Condition, F_2,32_=.06, p=.944; Group x Condition, F_2,32_=.15, p=.863).

**Fig. 5.**
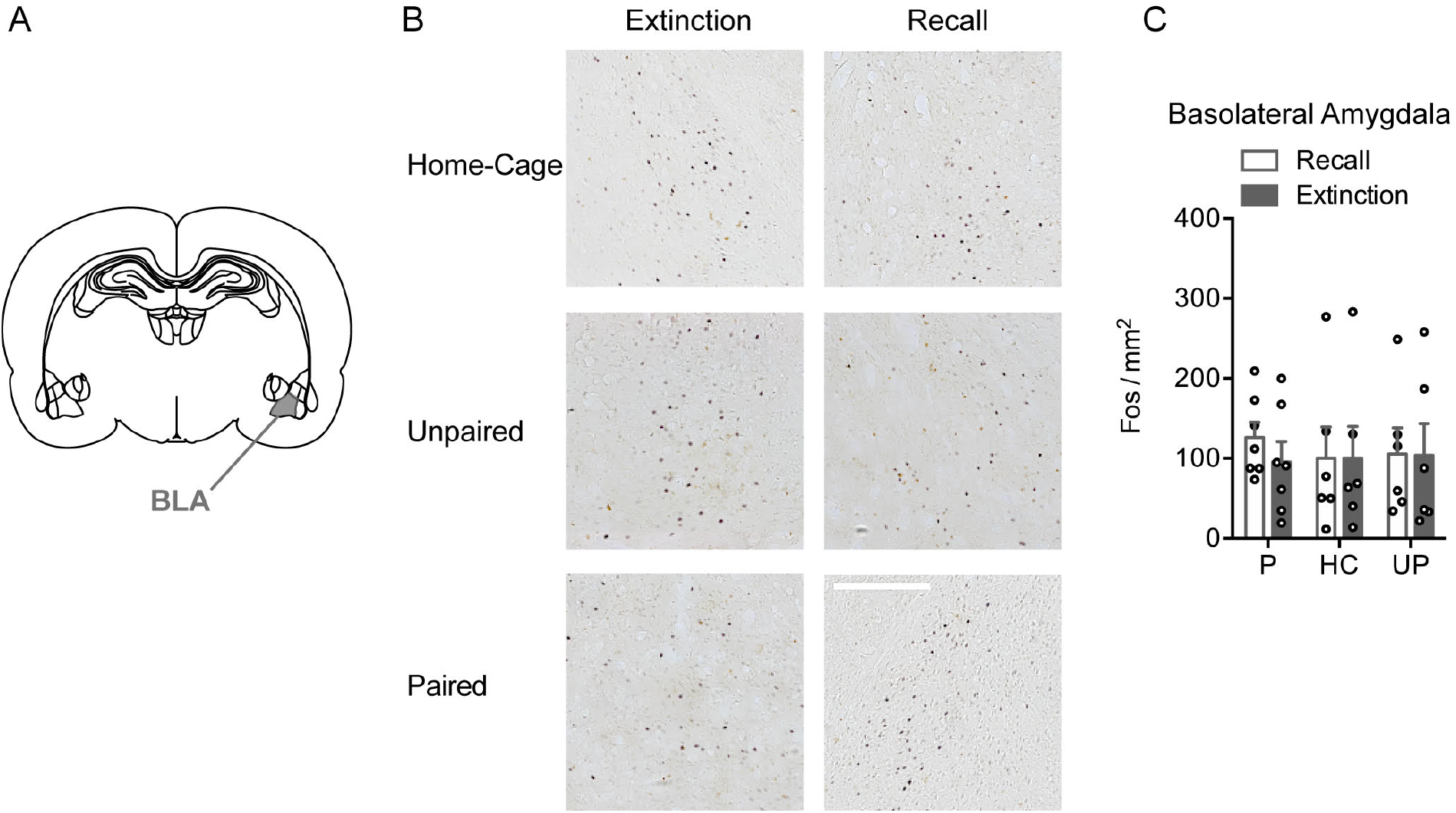
Fos density in the basolateral amygdala (BLA). **A** A schematic diagram indicates region where Fos was quantified in the BLA. **B** Representative Fos photomicrographs in the BLA. Scale bar is 250 μm. **C** Fos density in the BLA was not different between recall and extinction, and was similar between the paired, unpaired, and home-cage conditions.

#### Paraventricular nucleus of the thalamus

Fos density in the anterior and middle paraventricular nucleus of the thalamus (aPVT and mPVT) in the recall groups was found to be much greater in the paired condition relative to the unpaired and home-cage conditions, but this effect was not observed in the extinction groups. In the aPVT, Fos density was greater for the paired recall group compared to the paired extinction group (p=.006). Moreover, Fos density was greater for the paired recall group compared to the unpaired recall (p=.004) and home-cage recall groups (p=.003). All other post-hoc comparisons were not significant (p>.05) (Fig. 6A, B, C; Group, F_1,32_=1.09, p=.304; Condition, F_2,32_=5.51, p=.009; Group x Condition, F_2,32_=3.99, p=.028). Similarly, in the mPVT, Fos density was greater for the paired recall group compared to the paired extinction group (p=.003). Further, Fos density was greater for the paired recall group relative to the unpaired recall (p=.002) and home-cage recall groups (p<.001). No other statistical post-hoc comparisons were significant (p>.05) (Fig. 6A, D; Group, F_1,32_=2.56, p=.119; Condition, F_2,32_=8.61, p=.001; Group x Condition, F_2,32_=3.57, p=.040).

**Fig. 6.**
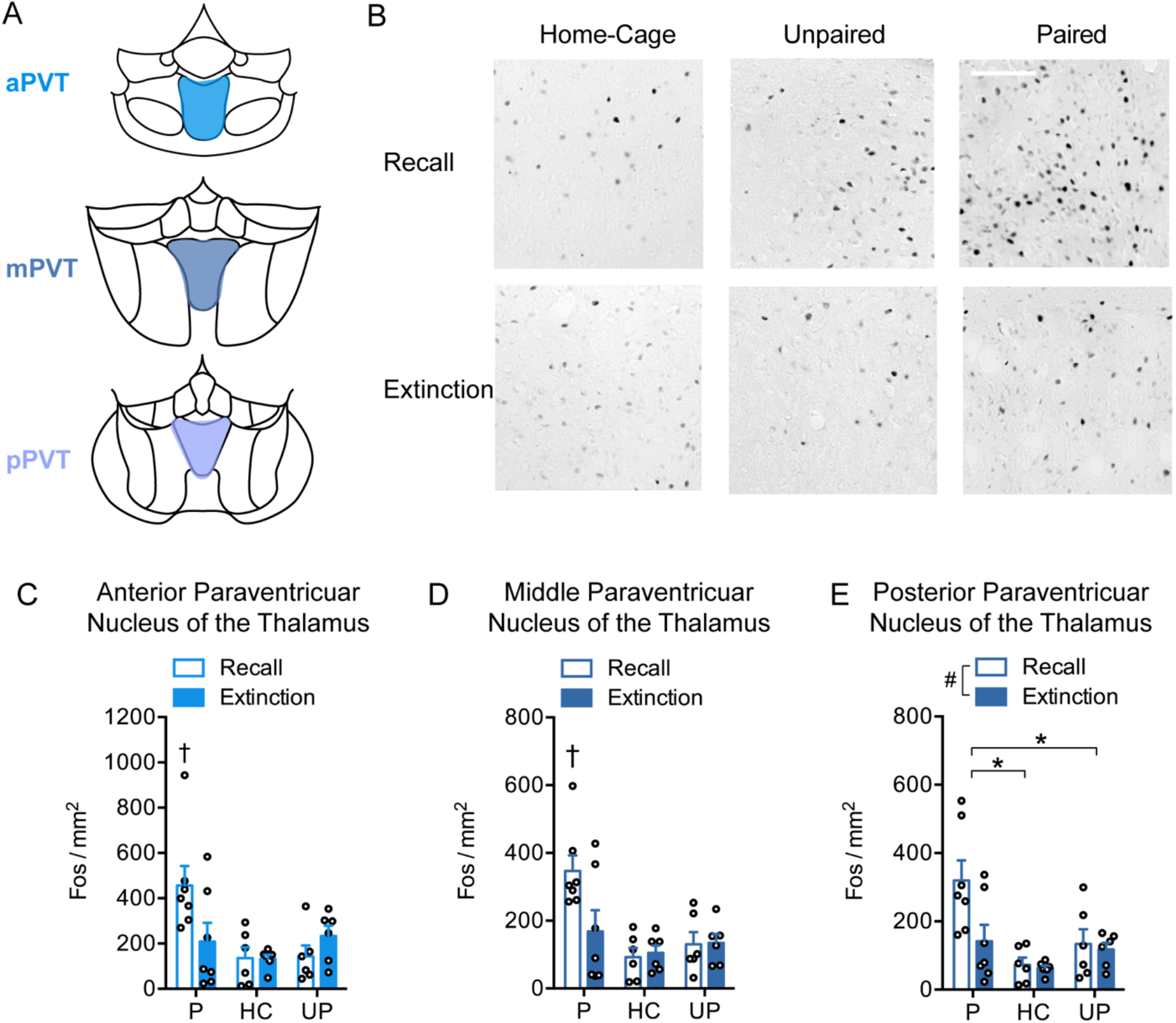
Fos density in the anterior paraventricular nucleus of the thalamus (aPVT), middle paraventricular nucleus of the thalamus (mPVT), and posterior paraventricular nucleus of the thalamus (pPVT). **A** Schematic diagrams indicating regions where Fos was quantified in the aPVT, mPVT and pPVT. **B** Representative Fos photomicrographs in the aPVT. Scale bar is 100 μm. **C** Fos density in the aPVT, and **D** mPVT, was greatest for the paired recall group compared to the paired extinction, unpaired recall and home-cage recall groups. **E** Fos density in the pPVT was greater following recall relative to extinction, and for the paired group relative to the unpaired and home-cage groups. †p<.05, significant group x condition interaction. #p<.05, recall vs extinction main effect. *p<.05, post hoc comparisons following main effect of training condition.

A similar pattern of results occurred in the posterior paraventricular nucleus of the thalamus (pPVT), but the interaction between Group and Condition did not reach statistical significance (Fig. 6A, E; Group x Condition, F_2,32_=3.00, p=.064). Fos density was greater during recall compared to extinction (Fig. 6F; Group, F_1,32_=4.36, p=.045; Condition, F_2,32_=8.96, p=.001). Fos density was greater for the paired condition compared to the unpaired (p=.036) and home-cage conditions (p=.001) but was not different between the unpaired and home-cage conditions (p=.493).

#### Correlational network analysis

To investigate the functional co-activation of the regions of interest, we computed correlations for each pair of regions within each group of animals. We then created inter-regional correlation matrices and network activation graphs for each experimental group following recall (Fig. 7A, B) and extinction (Fig. 8A, B). Network activation graphs display only strong and significant correlations (r≥.5, p<.05). The number of inter-regional correlations was greater for the paired extinction group compared to the paired recall group, with sparse connectivity in the home-cage and unpaired control conditions. Cortical sites showed an increase in connectivity during extinction and less so in recall. Moreover, we observed high correlations within related areas such as the different PVT subregions, and between the IL and PL, which were amongst the most widely correlated structures. Moreover, within the different PVT subregions, we observed increased node sizes in the paired recall, but not unpaired recall network activation graphs, reflecting an increase in Fos density compared to the home-cage controls. Interestingly, we found paired extinction specific network correlations with activity correlations between the IL and the mPVT, and between the mNAcSh and the aPVT and mPVT that were absent in the other groups.

**Fig. 7.**
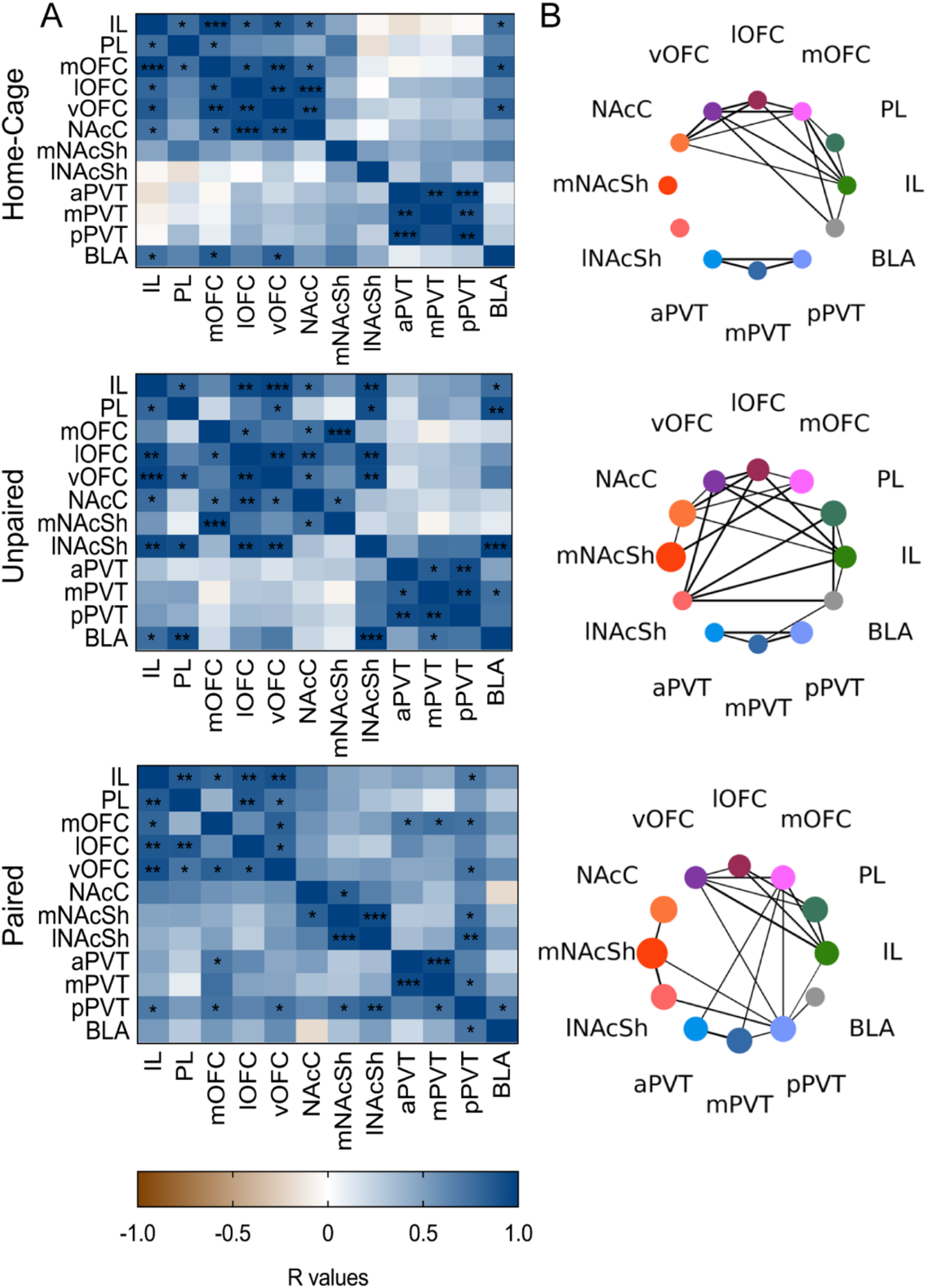
Cross-correlation and network activity analysis of Fos density in the home-cage (top panel), unpaired (middle panel), and paired (bottom panel) conditions during recall. **A** Pearson correlation matrices showing inter-regional correlations for Fos density. Axes represent brain regions. Colours reflect Pearson correlation coefficients and labels within squares correspond to P values of correlations. *p<.05; **p<.01; ***p<.001. **B** Network activation graphs indicate the strongest correlations (r≥.5, p<.05). Connecting line transparency represents correlation strength. Node size is proportional to the change of Fos density relative to the home-cage control condition.

**Fig. 8.**
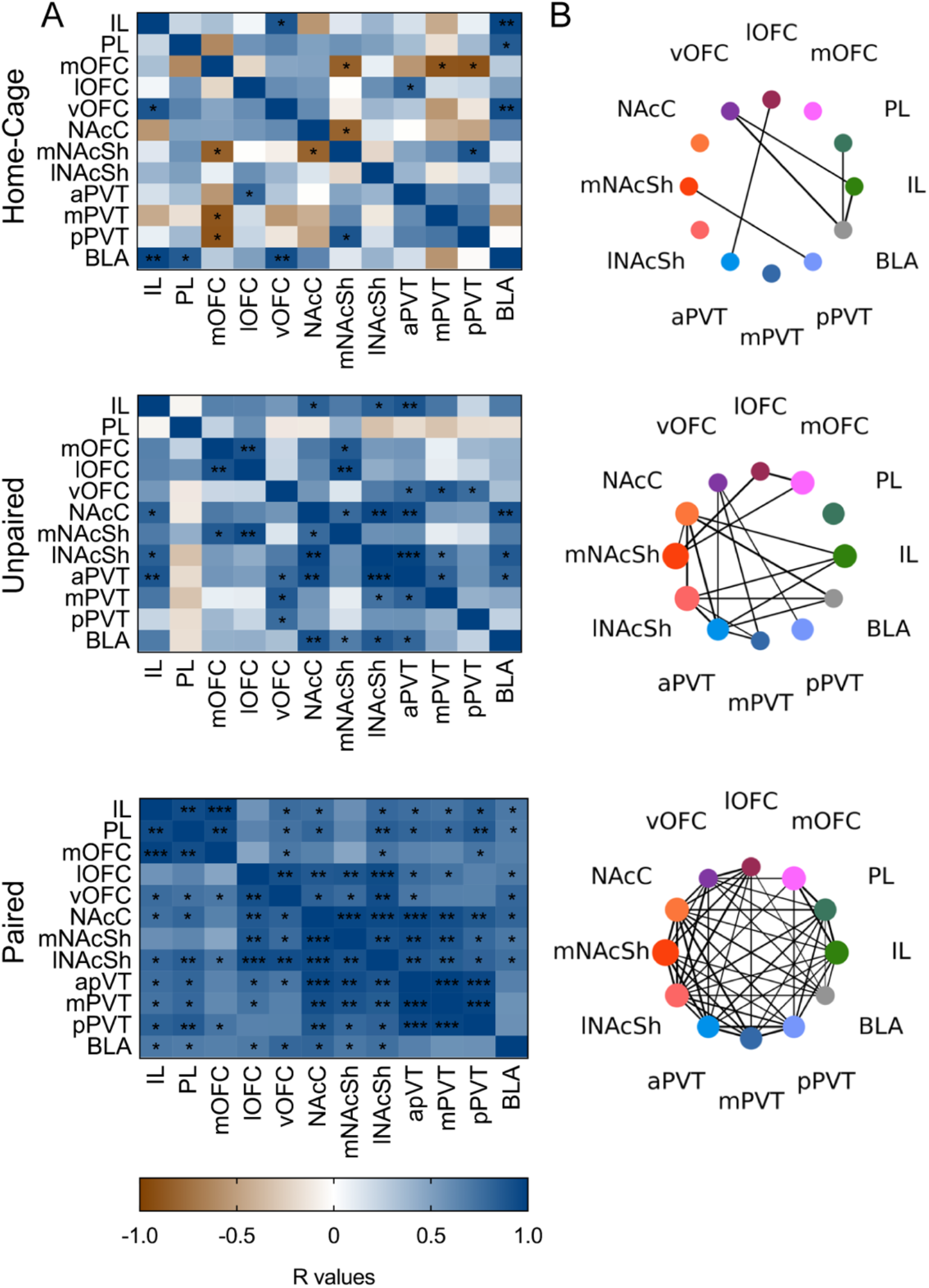
Cross-correlation and network activity analysis of Fos density in the home-cage (top panel), unpaired (middle panel), and paired (bottom panel) conditions during extinction. **A** Pearson correlation matrices showing inter-regional correlations for Fos density. Axes represent brain regions. Colours reflect Pearson correlation coefficients and labels within squares correspond to P values of correlations. *p<.05; **p<.01; ***p<.001. **B** Network activation graphs indicate the strongest correlations (r≥.5, p<.05). Connecting line transparency represents correlation strength. Node size is proportional to the change of Fos density relative to the home-cage control condition.

## Discussion

The present study investigated the neural correlates of appetitive Pavlovian extinction learning using Fos immunohistochemistry and correlational network connectivity analysis. We established a behavioural paradigm that allowed for the observation of differential Fos expression induced by the recall versus the extinction of responding to an appetitive Pavlovian CS. In most brain areas investigated here, we found greater Fos density for the paired and unpaired conditions relative to the home-cage condition, and no difference between the paired and unpaired conditions. However, in the BLA, vOFC, and lOFC, there were no differences between the paired, unpaired, and home-cage conditions. Fos density was only greater for the paired condition relative to the unpaired condition in the aPVT, mPVT and pPVT, suggesting that these regions are specifically recruited during responding elicited by the CS. Fos density was similarly elevated for the recall and extinction conditions in the IL, mOFC, mNAcSh and lNAcSh. In contrast, we found greater Fos density during recall compared to extinction in the PL, lOFC, NAcC, aPVT, mPVT, and pPVT. Lastly, we observed high correlations of Fos density across brain regions of interest especially in the paired extinction group compared to other groups. Together, our results provide novel network activation graphs of extinction and recall using an appetitive Pavlovian conditioning procedure and emphasise the role of the PVT in responding to discrete appetitive Pavlovian cues.

The neural correlates of extinction have been predominantly studied using Pavlovian fear conditioning and operant reward-seeking models (Hamlin et al., 2009; Perry & McNally, 2013; Warren et al., 2016; Bouton et al., 2021) as opposed to the appetitive Pavlovian learning design used here. During conditioning, rats in the paired groups showed an increase in ΔCS port entries across sessions indicating that they learned to associate sucrose reward with the CS, and rats in the unpaired and home-cage groups did not acquire a conditioned response. After one extinction session, the paired recall and the paired extinction groups displayed greater ΔCS port entries compared to the control groups, and after six extinction sessions, ΔCS port entries were reduced in the paired extinction group, while responding in the unpaired and home-cage groups remained low. Therefore, we established a behavioural paradigm that could efficiently condition and extinguish responding to a Pavlovian CS in the paired groups, and in which brain activation induced by recall of the CS on the first extinction session could be compared to brain activation induced by the CS following extinction.

In most brain regions examined here except for the PVT, Fos densities were increased above the home-cage conditions in both the paired and unpaired conditions. This finding was particularly evident in the IL and the PL. Further, we did not observe differences in Fos density between the paired and unpaired conditions in many brain regions. This suggests that many of these structures may be similarly recruited in both conditions. One argument could be that rats in the unpaired condition learned an association between sucrose delivery and the contextual stimuli present during training (i.e., a context-US association; Repucci & Petrovich, 2012). Therefore, brain regions may have been activated both by responding elicited by the CS (paired condition) and by the appetitive context (unpaired condition) (Repucci & Petrovich, 2012). Consistent with this idea, both the IL and PL integrate contextual information to guide behavioural responses (Hyman et al., 2012; Moorman & Aston-Jones, 2015). Similar results have been found using an appetitive Pavlovian conditioning task with a food reinforcer where Fos expression in the IL, PL and OFC were no different for paired and unpaired conditions (Yager et al., 2015). Therefore, similar Fos densities in the paired and unpaired conditions in other brain areas as well as the IL and PL may be due to different contextual associations that induce levels of neural activation similar to that induced by the CS-US association.

### Thalamic nuclei

Our results indicate that PVT neurons were selectively activated by the recall of appetitive Pavlovian conditioned responding, and showed reduced, nonselective responding to the CS following extinction. Subregions across the anterior-posterior axis of the PVT are composed of distinct neuronal subtypes (Gao et al., 2020) and are thought to have separate roles in reward learning (Barson et al., 2020). We found that Fos density in the aPVT and mPVT was significantly greater for the paired recall group vs the unpaired recall group. Although we found correlations in Fos density between all PVT subregions, this greater Fos density in the paired vs unpaired recall groups did not reach significance in the pPVT. It has similarly been reported that neurons in the aPVT, but not the pPVT, regulate operant sucrose-seeking during recall (Do-Monte et al., 2017). Interestingly, in all subregions of the PVT, Fos densities were greater for the paired condition relative to the unpaired condition, suggesting that this region is specifically recruited during responding to an appetitive Pavlovian CS. The PVT has been previously shown to be activated by cues that predict rewarding conditions, and by incentive stimuli previously paired with reward (Igelstrom et al., 2010; Flagel et al., 2011; Yager et al., 2015; Haight et al., 2017). Moreover, an incentive stimulus previously paired with a food reward has been shown to active cells in the PVT in a paired, but not unpaired, training condition (Yager et al., 2015). Therefore, the PVT may be specifically recruited by Pavlovian cues that predict reward.

Network analysis of Fos density showed network correlations that were specific to the paired extinction group with high activity correlations between the IL and mPVT, and between the mNAcSh and the aPVT and mPVT (Fig. 8B). This is consistent with dense glutamatergic projections from the IL to the PVT, and with strong projections from the PVT to the mNAcSh (Vertes, 2003; Vertes & Hoover, 2008). Moreover, inactivation of the IL-to-PVT pathway disrupts fear extinction retrieval, suggesting a role of IL inputs to the PVT in extinction (Tao et al., 2021). Further, the PVT-to-NAc pathway is involved in positive reinforcement and operant reward-seeking (Hamlin et al., 2009; Lafferty et al., 2020; Do-Monte et al., 2017). Together, our results suggest that the PVT is recruited by the recall of appetitive Pavlovian conditioned responding, and that Fos densities in the PVT are also correlated with neural activity in the IL and mNAcSh during extinction. These findings are consistent with the notion that cortical-striatal-thalamic connections involving the IL, mNAcSh, and PVT are important for appetitive extinction (Millan et al., 2011; McNally, 2014; Bouton et al., 2021).

### Cortical areas

We found greater Fos density in the PL during recall relative to extinction, which is consistent with a popular hypothesis that the PL drives the expression of freezing in Pavlovian fear conditioning and reward-seeking in appetitive operant conditioning (Peters et al., 2009). Support for this hypothesis comes from studies showing that PL inactivation disrupts operant reward-seeking, and attenuates renewal and reinstatement (Corbit & Balleine, 2003; McLaughlin & See, 2003; Fuchs et al., 2005; Trask et al., 2017). Consistently, inactivation of the PL disrupts the expression of conditioned fear but does not affect extinction learning or retrieval (Laurent & Westbrook, 2009; Choi et al., 2010; Sierra-Mercado et al., 2011). Similar to our results, Fos expression in the PL is greater during renewal as compared to extinction of responding to a Pavlovian CS paired with food (Anderson & Petrovich, 2017), and further, the PL does not express increased Fos during the extinction of operant food-seeking (Warren et al., 2016). Together with the current study, these results support the idea that the PL contributes to the expression rather than to the extinction of appetitive Pavlovian conditioned responding.

In the IL, we found similar expression of Fos density during recall and extinction. Since the recall session also functions as a first extinction session, one interpretation is that neural activity in the IL is maintained throughout early and extensive extinction learning. The IL has been shown to be active during early extinction of operant food-seeking (Warren et al., 2016), and others have also shown that neural activity in the IL is similar during both renewal and extinction of operant reward-seeking (Perry & McNally, 2013). This suggests that the IL may continue to function to consolidate extinction memories. Moreover, activation of the IL attenuates renewal, while IL lesions enhance the reinstatement of extinguished appetitive Pavlovian conditioned responding (Rhodes & Killcross, 2004; Villaruel et al., 2018). Similarly, IL inactivation disinhibits operant reward-seeking, promotes reinstatement, and disrupts the retrieval of extinction memories (Peters et al., 2008; Gutman et al., 2017). These results suggest that the IL functions to inhibit both operant and Pavlovian responding during tests of extinction retrieval.

A second interpretation of our results is that the IL has dual functions in both the recall and the extinction of appetitive Pavlovian conditioned responding. In support of this idea, mounting evidence suggests that discrete neural ensembles within the IL modulate opposing Pavlovian conditioned behaviours (Suto et al., 2016), and encode either operant reward-seeking or extinction memories (Warren et al., 2016; 2019). Thus, discrete neural ensembles within the IL may modulate the expression and the extinction of appetitive Pavlovian conditioned responding. This question could be addressed by determining whether the same neuronal ensembles remain active from recall to extinction, and whether recall and extinction induces activation in overlapping or distinct neuronal ensembles (Josselyn & Tonegawa, 2020).

Our results suggest that the mOFC and IL may have similar functional roles in appetitive Pavlovian recall and extinction, and that the ventral and lateral OFC play lesser roles. Like the IL, Fos density in the mOFC was comparable during recall and extinction. The mOFC and IL also respond similarly to Pavlovian cues that predict sucrose in decision-making and behavioural inhibition tasks (Bradfield et al., 2015; Hardung et al., 2017; Verharen et al., 2020). Fos density in the lOFC was greater following recall than extinction, although was not different for the paired, unpaired and home-cage controls. In the vOFC, we found no difference in Fos density between conditions or as a function of recall vs extinction, suggesting that in the vOFC, Fos density was not associated with recall or extinction. Inactivation of the lOFC, but not the mOFC, has been found to impair the reinstatement of operant reward-seeking (Fuchs et al., 2004; Arinze & Moorman, 2020). Moreover, inactivation of the lOFC has been shown to impair learning in a Pavlovian over-expectation task (Takahashi et al., 2009). However, our results are consistent with findings that vOFC and lOFC inactivation have no effect on the acquisition or extinction of conditioned responding to a Pavlovian CS that predicts sucrose (Burke et al., 2008, 2009). These findings are consistent with functional heterogeneity between distinct OFC subregions (Heilbronner et al., 2016). Together, our results suggest that the mOFC, but not the vOFC or lOFC, is involved in both the recall and extinction of responding to a Pavlovian conditioned appetitive CS.

### Striatal structures

The differential pattern of neural activity observed in the core and shell subregions of the NAc is consistent with evidence that these structures play distinct functional roles in learning and memory. In the NAcC, we found greater Fos density during recall relative to extinction, while in the mNAcSh and lNAcSh Fos density was similar during recall and extinction. Similarly, Fos expression in the NAcC is greater during renewal compared to extinction in an operant reward-seeking task, whereas Fos expression in the mNAcSh and lNAcSh is similar in renewal and extinction (Perry & McNally, 2013). Moreover, inactivation of the mNAcSh drives the reinstatement of operant drug-seeking (Peters et al., 2008). These results suggest that the NAcC promotes, and the mNAcSh suppresses, conditioned responding after extinction.

The IL-to-NAcSh pathway is thought to mediate the extinction of operant reward-seeking (Peters et al., 2009; Augur et al., 2016). However, we did not find correlated Fos expression in the IL and mNAcSh during extinction, suggesting that these areas have distinct activity patterns during extinction of appetitive Pavlovian conditioned responding. Recent evidence from our laboratory also suggests that the IL-to-mNAcSh pathway suppresses renewal, but not through an extinction mechanism, using an appetitive Pavlovian conditioning procedure (Villaruel et al., 2022). Therefore, the absence of correlated Fos expression in the IL and mNAcSh may be due to the use of an appetitive Pavlovian learning design.

### Amygdala

The BLA has been shown to have a prominent role in extinction learning using aversive Pavlovian conditioning (Laurent & Westbrook, 2008; Knapska and Maren, 2009; Zimmerman and Maren, 2010; Lingawi et al., 2019) and operant reward-seeking procedures (Fuchs & See, 2002; Fuchs et al., 2005; McLaughlin and See, 2003; McLaughlin & Floresco, 2007). Furthermore, others have shown using single-unit recordings that the BLA is involved in learning during over-expectation, although inactivation of the BLA has no effect on responding in over-expectation (Haney et al., 2010; Lucantonio et al., 2015). Therefore, even though we did not find that Fos density in the BLA was associated with the recall or extinction of appetitive Pavlovian responding, it may be that the technique used here was unable to detect the BLA’s involvement in these learning processes.

### Conclusions

We analyzed Fos density following either the recall of a CS-US association, or the extinction of responding to the CS, to evaluate the roles of multiple brain structures in recall and extinction of appetitive Pavlovian conditioned responding. We have found that the IL, mOFC, mNAcSh and lNAcSh showed increased activation following both recall and extinction vs the home-cage condition, but that activation was greater during recall vs extinction in the PL, NAcC and PVT, consistent with a selective role of these structures in promoting the expression of conditioned responding. The PVT showed enhanced activation in the paired condition relative to the unpaired condition during recall, suggesting that the PVT plays a special role in responding elicited by a discrete Pavlovian CS. However, most of the other brain areas examined here responded similarly in the paired and unpaired conditions, suggesting that they are similarly responsive to contextual cues that precede reward delivery and the CS. The present study provides novel evidence on the neural correlates underlying appetitive Pavlovian recall and extinction in multiple brain regions, and is consistent with the role of the PVT, and its connections with the IL and mNAcSh in controlling responding to an appetitive Pavlovian cue.

## Acknowledgements

This research was funded by a grant from the Natural Sciences and Engineering Research Council of Canada (NSERC) to N.C. N. C. was the recipient of a Chercheur-Boursier award from the Fonds de Recherche du Québec en Santé and a member of the Center for Studies in Behavioral Neurobiology. A.B. is supported by a doctoral scholarship from the Fonds de Recherche du Quebec en Nature et Technologies. F.R.V. is supported by a doctoral scholarship from NSERC. The authors would like to thank Melissa Martins, Ingrid Matei and William Edmiston for assistance with histology. The authors would also like to thank Steve Cabilio for technical support, Dr. Chris Law of the Center for Microscopy and Cellular Imaging for technical support and for writing the FIJI macro for Fos quantification, and Dr. Andrew Chapman for providing comments on the manuscript.

